# Antiviral reverse transcriptase–primase synthesizes protein-templated DNA

**DOI:** 10.64898/2026.04.18.719102

**Authors:** Xin-Yi Song, Yushan Xia, Jun-Tao Zhang, Xin-Yang Wei, Hua Qi, Linshu Li, Ning Jia

## Abstract

Defense-associated reverse transcriptases (DRTs) are widespread in bacteria^1,2^, but how multi-domain DRTs containing RT and additional catalytic activities coordinate antiviral defense remains unclear. Here we show that DRT7, which contains both reverse transcriptase (RT) and primase–polymerase (PP) domains, provides broad-spectrum anti-phage immunity through abortive infection and can be activated by a phage-encoded putative transcriptional regulator. Upon activation, DRT7 synthesizes long, protein-primed, palindromic poly(A)/poly(T)-rich duplex-like DNA. Cryo–electron microscopy structures reveal that RT initiates protein-primed, protein-templated, sequence-specific poly(T) synthesis through an arginine-rich recognition pocket without requiring a complementary nucleic acid template, thereby converting DRT7 from an inactive closed dimer to an active open dimer. The RT-produced poly(T) then serves as both primer and template for PP-mediated poly(A) extension, with iterative handoff between RT and PP generating palindromic, alternating poly(A)/poly(T) ssDNA tracts that assemble into fold-back duplex-like DNA. These findings uncover an unexpected antiviral strategy based on protein self-templating, sequence-specific duplex-like DNA synthesis and reveal how coupling RTs with additional catalytic activities expands the functional scope of nucleic acid synthesis pathways.

## Main

Reverse transcriptases (RTs) are RNA-dependent DNA polymerases that convert RNA into complementary DNA (cDNA). First discovered in retroviruses in 1970^3,4^, RTs became important tools in molecular biology^5–8^ and are now recognized as widespread across all domains of life^1,2,9,10^. In bacteria, RTs were identified as components of retrons^11–14^ and only recently recongnized for their role in antiviral (anti-phage in bacteria) defense^2,15^. Multiple RT-based anti-phage systems have now been identified, including CRISPR-associated RTs, abortive infection (Abi) RTs, retrons, and diverse unknown groups (UGs)^1,2,9^.

UG RTs exhibit substantial diversity and are frequently associated with noncoding RNAs or additional protein domains involved in anti-phage defense^1,16^. Early examples include the abortive infection proteins AbiK, AbiA, and Abi-P2^17–19^, which restrict phage replication through protein-primed, template-independent DNA synthesis^20,21^. More recent analyses identified a subset of UG RTs termed defense-associated RTs (DRTs)^1^. Some function as standalone RTs that generate untemplated DNA such as DRT4^22^, whereas others rely on RNA templates to produce antiviral single-stranded or double-stranded DNA, such as DRT2^23,24^, DRT9^25,26^ and DRT10^27^. Whereas these characterized RT defense systems involve standalone RTs or RT–ncRNA complexes, DRTs lacking ncRNA but containing additional catalytic domains remain poorly understood. One such system, DRT7, contains a reverse transcriptase domain fused to a primase–polymerase (PP) domain, both required for anti-phage activity^1^. Similar RT-PP fusions have been identified in other defense systems, including CRISPR–Cas–associated RTs^28^ and DRT1 systems^2^. However, the molecular mechanisms underlying how RT and PP are coordinated within a single protein to mediate anti-phage defense remain unknown.

Here, we uncover a novel antiviral RT- and PP-based immunity in which iterative cooperation between reverse transcriptase and primase–polymerase activities generates long, protein-primed, protein-templated, sequence-specific, palindromic poly(A)/poly(T)-rich duplex-like DNA. This mechanism illustrates how coupling reverse transcriptase activity with primase–polymerase function enables de novo protein-templated DNA synthesis and expands the functional scope of nucleic acid synthesis pathways.

### DRT7 confers broad-spectrum anti-phage defense via abortive infection

To elucidate the mechanism of DRT7-mediated anti-phage defense, we cloned the *drt7* gene from *Escherichia coli* strain SC366, together with its native promoter, into *E. coli* DH10B (Fig. 1a). Expression of DRT7 conferred broad-spectrum resistance to bacteriophages from the BASEL collection, with up to ∼10^6^-fold reductions in efficiency of plating (EOP) (Fig. 1b), consistent with previous observations^1^. DRT7 did not impair phage adsorption (Fig. 1c), but instead inhibited infection at a post-adsorption stage, preventing progeny release (Fig. 1d). Resistance to phage T4 was observed at low multiplicities of infection (MOI = 0.05 and 0.5) but was largely overcome at high MOI (MOI = 5), which induced growth arrest in infected cells, consistent with an abortive infection (Abi) defense strategy (Fig. 1e)^29^. Disruption of conserved catalytic residues in either the PP domain (D68A/D70A; DxD→AxA), the RT domain (D626A/D627A; YVDD→YVAA), or deletion of the C-terminal region (Δ666–1776) completely abolished anti-phage activity (Fig. 1f), demonstrating that all three regions are essential for DRT7 function.

**Figure 1.**
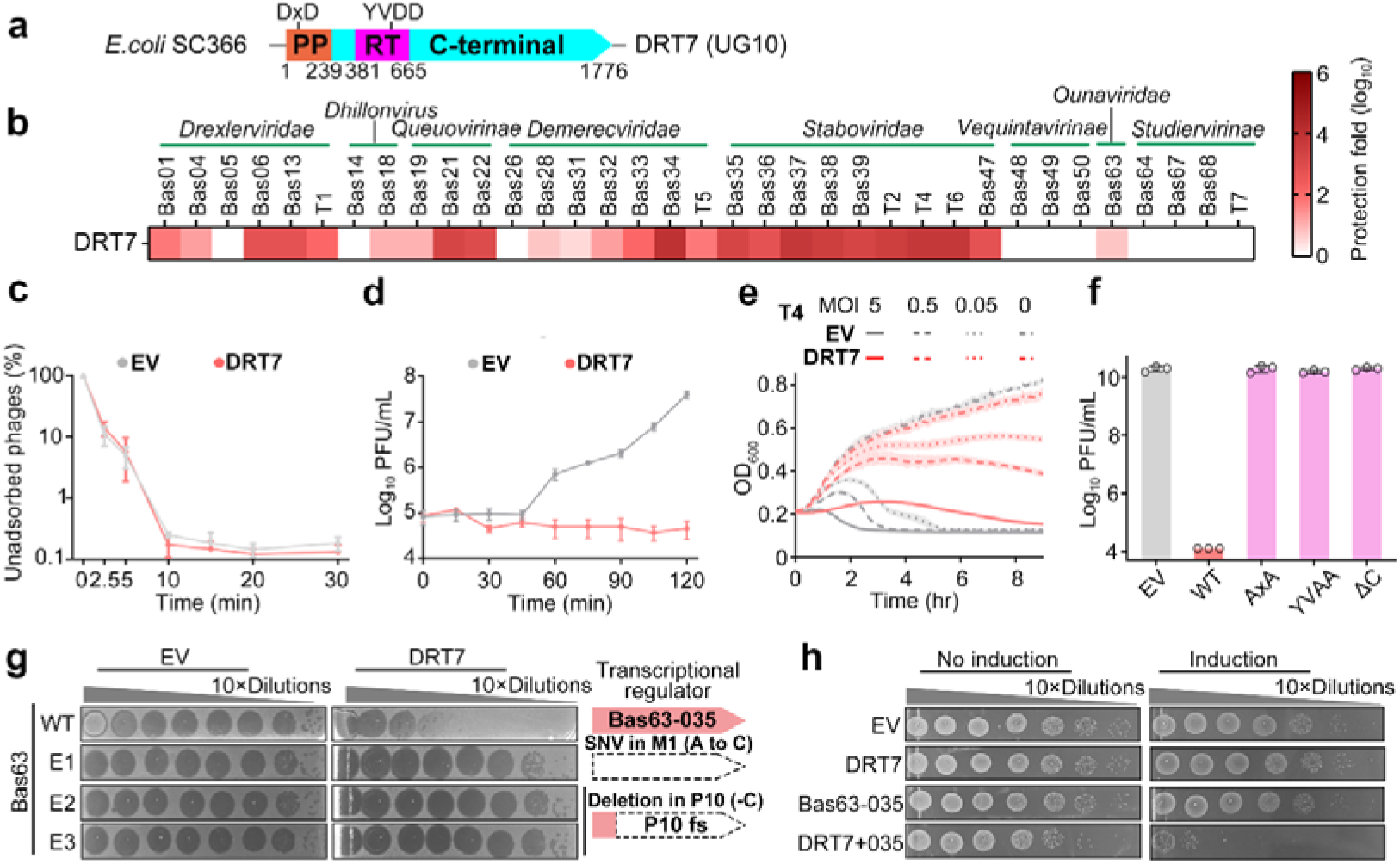
DRT7 restricts phage propagation through abortive infection. **a,** Schematic of the DRT7 from *Escherichia coli* strain SC366. PP, primase–polymerase; RT, reverse transcriptase. **b,** Plaque assays showing susceptibility of *E. coli* DH10B expressing DRT7 or an empty vector (EV) to bacteriophages from the BASEL collection. Representative images from three biological replicates are shown. **c**, Phage adsorption to cells expressing DRT7 or EV. **d**, One-step growth curves of phage T4 infecting DH10B cells with or without DRT7 expression. **e,** Growth of DH10B cells expressing DRT7 or EV following T4 infection at multiplicities of infection (MOI) of 0, 0.05, 0.5, or 5. **f,** Anti-phage activity of wild-type DRT7 and catalytic mutants. DH10B cells expressing wild-type DRT7, the PP AxA mutant (D68A/D70A; DxD→AxA), the RT YVAA mutant (D626A/D627A; YVDD→YVAA), the C-terminal truncation (Δ666–1776, ΔC), or EV were challenged with T4 phage. Data represent mean ± SD from three biological replicates, with individual data points shown. **g,** The collected plaques were tested for their ability to overcome the defense systems through a series of dilution plaque assays. Three Bas63 phage mutants were identified, each carrying mutations in putative transcriptional regulator Bas63-035. **h,** Survival status of *E. coli* cells co-producing DRT7 system and Bas63-035 protein.

Given that abortive infection systems are typically activated upon phage invasion, we next sought to identify factors that trigger DRT7 activation. To this end, we isolated phage mutants capable of escaping DRT7-mediated immunity. Challenge of E. coli DH10B cells expressing DRT7 with phage Bas63 yielded three independent escape mutants (Fig. 1g). Whole-genome sequencing revealed that all three mutants carried mutations in the putative transcriptional regulator gene Bas63-035, including start codon substitutions or frameshift deletion (Fig. 1g). Consistently, cell survival assays showed that cytotoxicity occurred only when both DRT7 and Bas63-035 were co-expressed (Fig. 1h), indicating that this phage-encoded factor activates the DRT7 system. Together, these results demonstrate that DRT7 restricts phage propagation through an abortive infection mechanism that requires coordinated RT and PP activities as well as the C-terminal region, and that activation of this defense is triggered by a phage-encoded transcriptional regulator.

### DRT7 catalyzes protein-primed synthesis of palindromic poly(A)/poly(T)-rich duplex-like DNA

To further investigate the molecular mechanism of DRT7-mediated anti-phage defense, we tested whether DRT7 can synthesize DNA, given it contains PP and RT domains (Fig. 1a). The results showed that DRT7 catalyzed DNA synthesis, with both product abundance and length increasing as dNTP concentrations increased (Fig. 2a). This activity required divalent metal ions, with both Mg²□ and Mn²□ supporting catalysis and Mg²□ being effective at physiological concentrations (Extended Data Fig. 1a). Addition of dNTPs caused an upward mobility shift of DRT7 on SDS–PAGE that was abolished by DNase treatment, indicating covalent attachment of newly synthesized DNA to the protein (Fig. 2b), reminiscent of protein-primed DNA synthesis similar to the AbiK and DRT9 defense systems^20,21,25,26^. Illumina sequencing of these products revealed strong enrichment of adenine- and thymine-rich sequences (>60% A or T), frequently containing ≥15-nt homopolymeric tracts (Fig. 2c). Consistent with *in vitro* observations, following phage infection, wild-type DRT7 cells exhibited higher levels of poly(A)- and poly(T)-rich reads than the catalytically inactive YVAA mutant, indicating activation of DRT7-dependent DNA synthesis for anti-phage defense *in vivo* (Extended Data Fig. 1b).

**Figure 2.**
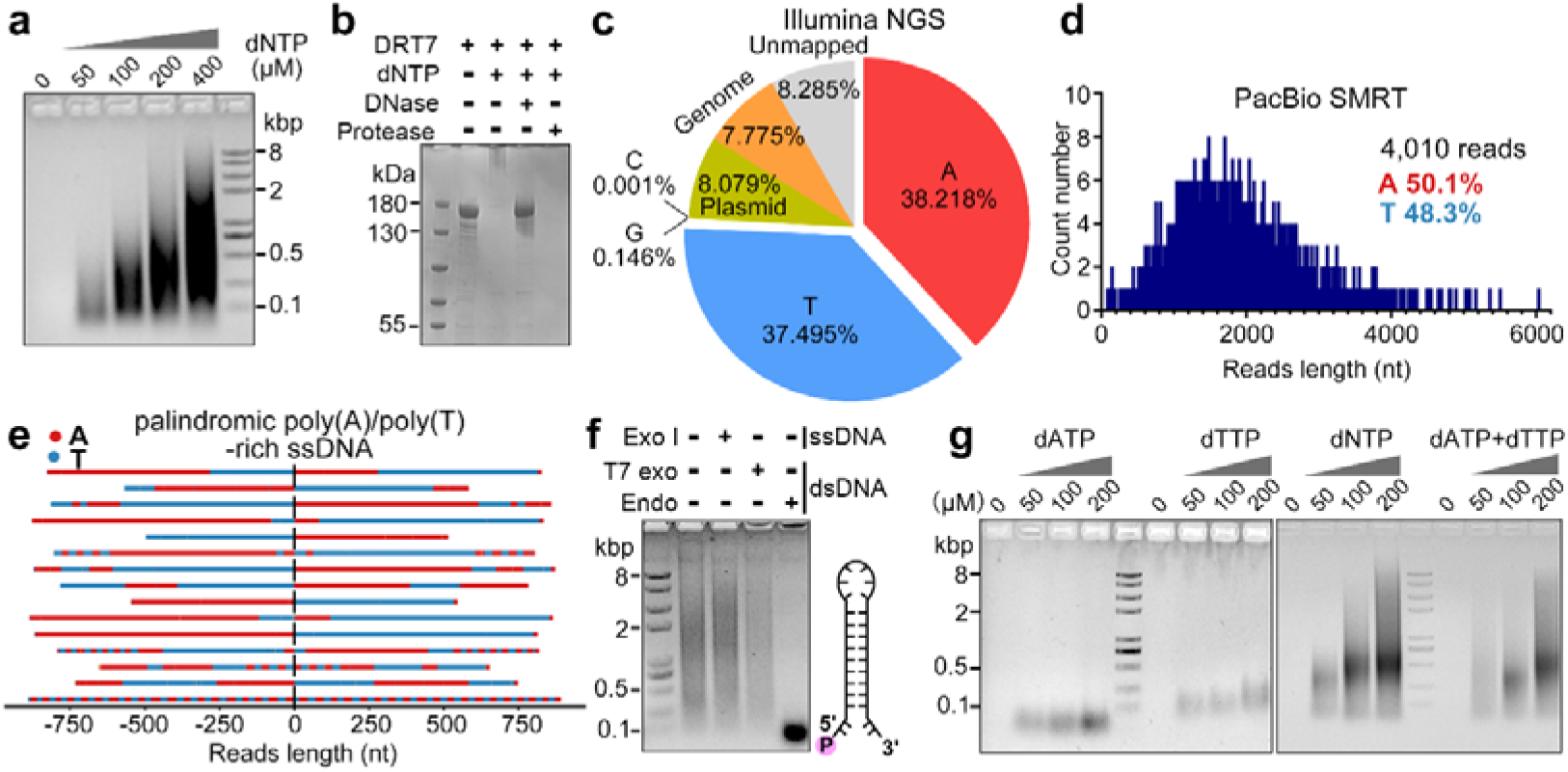
DRT7 synthesizes long, palindromic poly(A)/poly(T)-rich single-stranded DNA. **a,** Agarose gel analysis of DNA products generated by 400 nM DRT7 in reactions containing increasing dNTP concentrations. **b,** SDS–PAGE analysis of DRT7 incubated with or without dNTPs. dNTP addition induces an upward mobility shift, which is eliminated by DNase or protease treatment. **c,** Nucleotide composition of Illumina sequencing reads from *in vitro* DRT7 products, showing strong enrichment for adenine- and thymine-rich sequences. **d,** Length distribution of PacBio long reads, revealing predominantly A/T-rich DNA with an average length of ∼1.8 kb. **e,** Representative PacBio reads aligned at their centers to highlight palindromic symmetry. **f,** Nuclease digestion of DRT7 products. DNA is resistant to single-strand-specific exonuclease I and 5′→3′ double-strand–specific T7 exonuclease but is cleaved by double-strand–specific endonucleases, consistent with protein-linked, self-complementary ssDNA (schematic, right). **g,** Agarose gel analysis of DRT7 reactions containing dATP, dTTP, both nucleotides, or all four dNTPs, showing that dATP and dTTP alone are sufficient to support synthesis of long DNA products.

To determine the sequence composition of the synthesized DNA, we performed PacBio SMRT sequencing of the *in vitro* products, which revealed molecules composed almost exclusively of adenine (50.1%) and thymine (48.3%), with an average length of 1,812 nucleotides (Fig. 2d). Individual molecules consisted of alternating poly(A) and poly(T) tracts and displayed a palindromic organization predicted to form stable intramolecular hairpin structures (Fig. 2e). These DNA products were resistant to single-strand–specific Exonuclease I and double-strand–specific T7 exonuclease but were cleaved by double-strand–specific endonucleases (Fig. 2f), indicating protein-primed synthesis of self-complementary ssDNA whose 5′ ends remain covalently attached to DRT7 (Fig. 2b, e). Consistent with this alternating poly(A)/poly(T) architecture, DRT7 failed to generate long DNA products when supplied with only dATP or dTTP, whereas robust synthesis occurred when both nucleotides were present, reaching levels comparable to reactions containing all four dNTPs (Fig. 2g). Together, these results demonstrate that DRT7 synthesizes protein-primed, palindromic, poly(A)/poly(T)-rich ssDNA that forms self-complementary intramolecular hairpin duplex-like structures for anti-phage defense.

### DRT7 adopts an inactive closed dimer bound to protein-primed, protein-templated poly(T) ssDNA

To elucidate how DRT7 generates palindromic ssDNA, we determined cryo–electron microscopy (cryo-EM) structures of DRT7 in the presence of dATP and dTTP (Fig. 3a-e and Extended Data Fig. 2, 3). Two-dimensional classification revealed two dimeric assemblies and a monomeric population (Extended Data Fig. 2a). Because size-exclusion chromatography and mass photometry indicated that DRT7 is predominantly dimeric in solution (Extended Data Fig. 4a, b), we attribute the monomeric particles to grid-induced dissociation and focused our analysis on the dimeric reconstructions. One class assembles into a closed dimer in which two near-identical protomers assemble in a back-to-back configuration, burying ∼1,000 Å² of surface area (Fig. 3b and Extended Data Fig. 2a). In each protomer, a 10-nt poly(T) oligonucleotide is observed within a channel formed by the RT and αRep domains (Fig. 3c). The electron density between positions −5 and −6 is discontinuous, suggesting heterogeneity in DNA length among protomers (Fig. 3c, f and Extended Data Fig. 3). The 5′ end of the poly(T) is covalently linked to Y682 in the αRep domain, providing direct structural evidence for protein-primed DNA synthesis (Fig. 3c, right inset). The absence of palindromic poly(A)/poly(T)-rich ssDNA in this conformation indicates that the closed dimer represents an inactive state.

**Figure 3.**
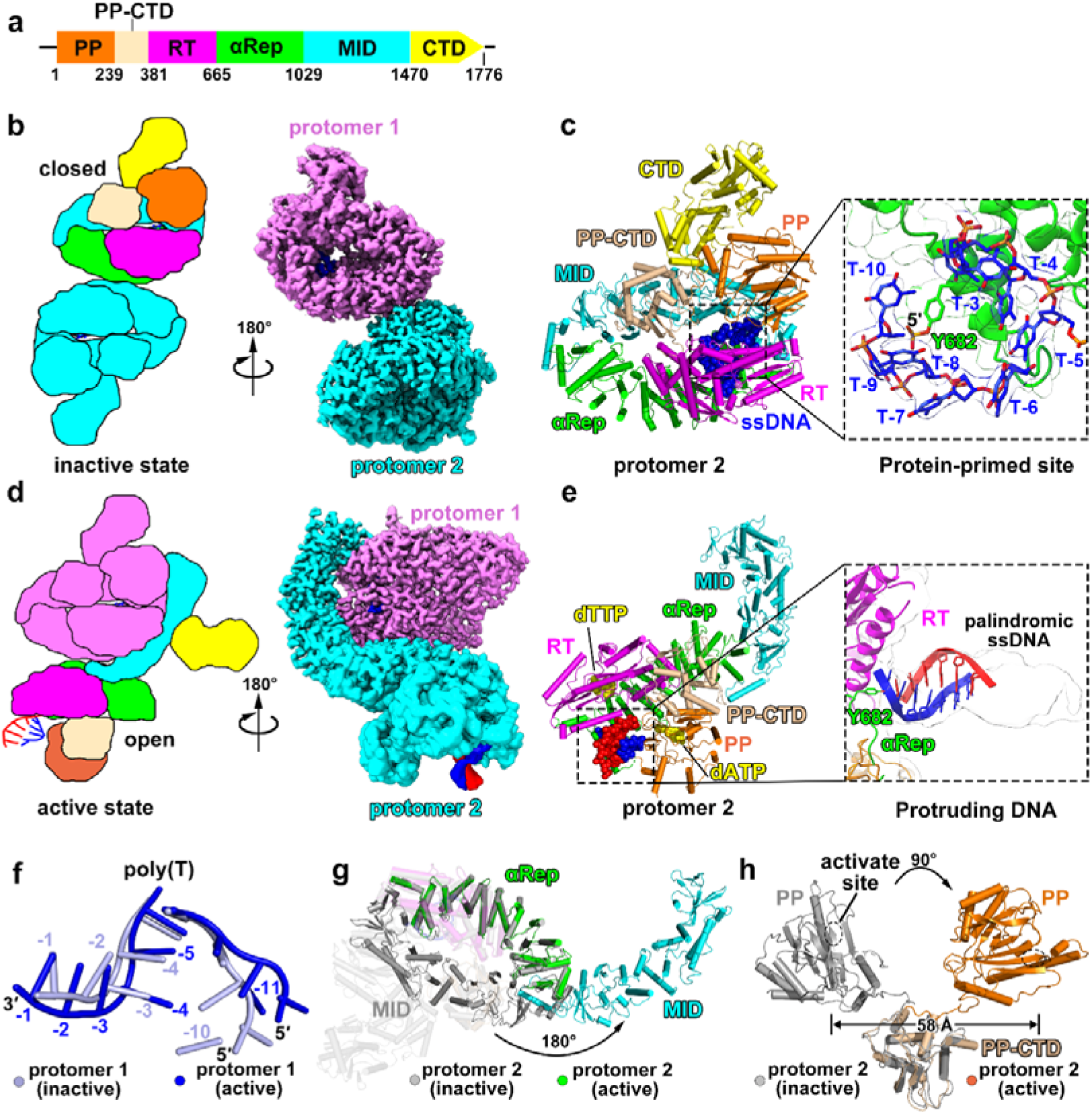
Structural basis of DRT7 activation from a closed inactive dimer to an open active dimer. **a,** Domain organization of DRT7. **b, d,** Schematic and surface representations of DRT7, showing the closed inactive dimer (**b**) and open active dimer (**d**). **c,** Cartoon representation of protomer 2 in the inactive state with a bound 10-nt poly(T) oligonucleotide. **e,** Cartoon representation of protomer 2 in the active state bound to a palindromic poly(A)/poly(T)-rich ssDNA product. **f,** Comparison of ssDNA bound to protomer 1 in the inactive and active states. **g,** Conformational transition between the closed (inactive, gray) and open (active, colored) states of protomer 2, involving an ∼180° rotation of the MID domain relative to the αRep domain. **h,** Superposition of the PP domains from protomer 2 in the inactive (gray) and active (wheat) states, highlighting large rotational rearrangements.

Structural analysis further revealed that, in addition to the PP (residues 1–239) and RT (residues 382–665) domains, DRT7 contains four additional modules: a PP C-terminal domain (PP-CTD; residues 240–381), an α-helical repeat domain (αRep; residues 666–1029), a middle domain (MID; residues 1030–1470), and a C-terminal domain (CTD; residues 1471–1776) (Fig. 3a and Extended Data Fig. 5a). The spatial arrangement of the RT and αRep domains resembles that of AbiK, a protein-primed reverse transcriptase involved in template-independent ssDNA synthesis (Extended Data Fig. 5b, c)^20^. The PP domain is structurally similar to CRISPR-associated primase–polymerases (CAPPs) encoded within CRISPR–Cas operons implicated in adaptation (Extended Data Fig. 5d)^30,31^, whereas the PP-CTD shows distant similarity to the C-terminal domain of eukaryotic DNA primase but lacks an iron–sulfur cluster (Extended Data Fig. 5e)^32^. Together, these data indicate that DRT7 adopts a dimeric architecture bound to protein-primed poly(T) synthesis, representing an inactive state.

### Elongation of poly(T) enables open dimer formation to activate DRT7

A second class from 3D classification adopts an open dimer in which protomer 1 remains in a closed conformation, whereas protomer 2 transitions to an open conformation that wraps around protomer 1, burying ∼2,700 Å² of surface area (Figure 3D). In this assembly, an 11-nt poly(T) was observed, one nucleotide longer than in the inactive dimer (Fig. 3f and Extended Data Fig. 5f, g). In contrast, protomer 2 contains a protruding protein-primed, palindromic poly(A)/poly(T)-rich ssDNA that is covalently attached to residue Y682 and a single dTTP molecule located in the active site (Fig. 3e), which represents the active state. Structural comparison revealed that protomer 1 remains largely unchanged between inactive and active assemblies (RMSD=1.6 Å), with the primary difference being extension of the RT-synthesized poly(dT) from 10 to 11 nucleotides (Fig. 3f and Extended Data Fig. 5f, g), contributing to stabilizing the open protomer 2. The protomer 2 undergoes extensive remodeling: the MID domain rotates by ∼180° relative to the αRep domain (Fig. 3g), generating an open configuration that engages protomer 1 through multiple interactions mediated by three distinct dimer interfaces involving the RT, αRep, and MID domains (Extended Data Fig. 6a, b). Mutation of residues in these interfaces (RT-αRep, R–α; αRep-MID, α–M; or MID-MID, M–M) abolishes DRT7-mediated anti-phage defense (Extended Data Fig. 6c), demonstrating the essential role of the open dimer assembly for anti-phage function. This rearrangement releases constraints on the PP and CTD domains, allowing the PP domain to reposition via an ∼90° rotation relative to the PP-CTD domain (Fig. 3h). Together, these findings indicate that DRT7 functions as an open active dimer, with protomer 1 serving as a closed structural scaffold and protomer 2 acting as an open catalytic subunit responsible for palindromic ssDNA synthesis.

### The RT domain initiates protein-primed, protein-templated, thymidine-specific poly(T) synthesis

Sequencing of DRT7 reaction products showed that DRT7 generates palindromic, poly(A)/poly(T)-rich ssDNA (Fig. 2e), and structural analysis revealed that these DNA products are covalently linked to Y682 in the active conformation (Fig. 3e). In addition, a poly(T) oligonucleotide with its 5′ end linked to Y682 and its 3′ end positioned in the RT catalytic pocket was observed in the inactive conformation (Fig. 3c). Together, these observations indicate that DNA synthesis is initiated by a Y682-primed poly(T) intermediate generated by the RT domain, which is arrested in the inactive state and subsequently extended in the active assembly to produce palindromic poly(A)/poly(T)-rich ssDNA. We next investigated how the RT domain achieves thymidine selectivity in the absence of an RNA template. Structural analysis of protomer 2 in the inactive state, which later converts to the active form and is responsible for palindromic poly(A)/poly(T)-rich duplex-like ssDNA synthesis, revealed key features of the RT active site following poly(T) elongation. In addition to the conserved YVDD catalytic motif (Y624-D627, D526) and Y531, which stabilizes the ribose moiety of the T-1 corresponding to the incoming dTTP, four arginine residues (R440, R447, R595, and R750) form a highly selective initiation pocket that recognizes the thymine base at −1 position (Fig. 4a). Specifically, R440 and R750 form hydrogen bonds with thymine carbonyl oxygen O4, R595 contributes additional base-specific hydrogen-bonding interactions with thymine carbonyl oxygen O2, and R447 engages in a cation–π interaction with the thymidine base. Together, these interactions restrict productive binding to dTTP over other dNTPs. Mutation of the YVDD catalytic motif, D526 or the base-specific residues R440, R447, or R595 abolished anti-phage activity *in vivo* and eliminated DNA synthesis *in vitro* (Fig. 4b, c), demonstrating that both RT serves as a template for dTTP selection, which are essential for DRT7 anti-phage function. Beyond the initiation site, downstream nucleotides within the poly(dT) tract are stabilized by sequence-specific interactions involving D685 (T-4), H725 (T−5), F494 and N485 (T−6), N1399, T1401 and S1435 (T-9), further highlighting thymidine selectivity within the RT domain (Extended Data Fig. 7a). Together, these results demonstrate that the RT domain of DRT7 initiates protein-primed, protein-templated, thymidine-specific poly(T) synthesis through an arginine-rich recognition pocket, and that this specificity is essential for DRT7-mediated anti-phage defense.

**Figure 4.**
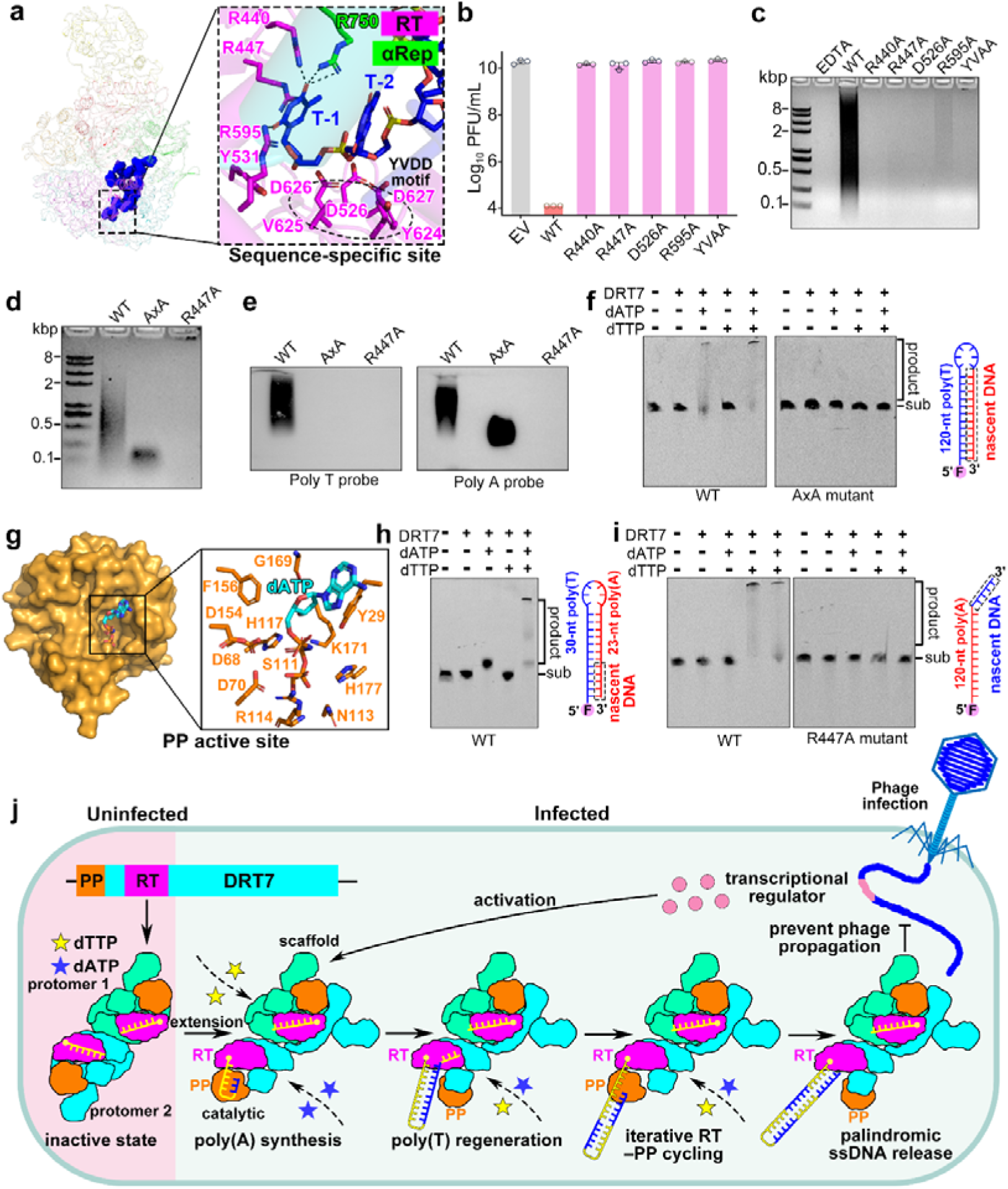
Iterative RT–PP handoff drives synthesis of palindromic poly(A)/poly(T)-rich ssDNA. **a,** Structural basis for thymidine-specific synthesis in the RT domain. **b,** Anti-phage activity of wild-type DRT7 and thymidine-recognition mutants in *E. coli* DH10B challenged with T4 phage. Data represent mean ± SD from three biological replicates. YVAA represents the RT catalytic mutant (D626A/D627A; YVDD→YVAA). **c,** Agarose gel analysis of DNA synthesis by 400 nM wild-type DRT7 and the indicated mutants incubated with 800 μM dATP and dTTP, with or without 10 mM EDTA. **d,** DNA products synthesized by wild-type DRT7, the PP-deficient AxA mutant (D68A/D70A; DxD→AxA), or the RT mutant (R447A). **e,** Southern blot detection of ssDNA using poly(A)- or poly(T)-specific probes following *in vitro* synthesis. **f,** Extension of a synthetic 5′-FAM–labeled 120-nt poly(T) substrate by wild-type DRT7 but not the PP mutant (AxA). **g,** Active sites of the PP domain bound to dATP. **h,** Extension of 5′-FAM–labeled hybrid poly(T)/poly(A) substrates. **i,** Extension of a synthetic 5′-FAM–labeled 120-nt poly(A) substrate by wild-type DRT7 but not the RT catalytic mutant (R447A). **j,** Proposed model for DRT7-mediated defense: phage-triggered activation promotes iterative RT–PP cycling, generating palindromic poly(A)/poly(T)-rich ssDNA that restricts phage propagation.

### RT-synthesized poly(T) primes PP-mediated poly(A) extension

Sequencing showed that poly(A) tracts are added immediately downstream of poly(T) (Fig. 2e), while the resulting palindromic ssDNA remains covalently linked to DRT7 (Fig. 3e), indicating that the RT-generated poly(T) functions as a primer for the following poly(A) extension. Because the RT active site is specialized for thymidine incorporation, we hypothesized that poly(A) synthesis is mediated by the PP domain. Mutation of the PP catalytic residues (D68A/D70A, DxD→AxA) abolished formation of long poly(A)/poly(T)-rich products, yielding only short poly(T) oligonucleotides (Fig. 4d), as confirmed by Southern blotting (Fig. 4e). These products remained covalently linked to DRT7, as shown by SDS–PAGE analysis (Extended Data Fig. 7b), indicating the initial poly(T) synthesis is mediated by the RT domain. Thus, PP inactivation selectively blocked downstream poly(A) extension while preserving RT-mediated poly(T) synthesis. By contrast, mutation of the RT residue R447 abolished both the initial poly(T) intermediate and the downstream poly(A)/poly(T)-rich products (Fig. 4d, e and Extended Data Fig. 7b), indicating that RT-generated poly(T) is required to initiate subsequent synthesis. These results support a sequential model in which the RT domain generates a protein-primed initial poly(T) primer that is extended by the PP domain through poly(A) synthesis.

To directly test whether the PP domain can extend a poly(T) primer with dATP, we supplied an exogenous 5′ FAM-labeled 120-nt poly(T) oligonucleotide and assayed dATP incorporation. DRT7 efficiently extended the poly(T) primer with dATP, whereas PP catalytic mutants abolished this activity (Fig. 4f). Consistently, SDS–PAGE analysis revealed a mobility shift of DRT7 upon addition of dATP, which was abolished by DNase treatment and was absent in PP catalytic mutants (Extended Data Fig. 7c), indicating that the PP domain catalyzes DNA synthesis on the RT-generated poly(T) primer. Together, these results reveal that RT-synthesized poly(T) serves as a protein-primed primer that is subsequently extended by the PP domain through poly(A) synthesis, establishing a sequential handoff from the RT to the PP domain.

### Poly(T) also serves as the template for PP-mediated poly(A) synthesis

We next asked what determines the dATP specificity of the PP domain. In the active state, a dATP molecule was observed bound in the PP active site (Fig. 4g). Notably, whereas the RT domain selects thymidine through extensive base-specific contacts (Fig. 4a), the PP active site lacks residues that directly recognize adenine and instead resembles the template-guided architecture of MsCAPP (Fig. 4g and Extended Data Fig. 7d, e). With dATP positioned opposite the template strand, nucleotide selection is likely governed by Watson–Crick base pairing, likely using the RT-synthesized poly(T) strand as the template. To directly test this hypothesis, we used a DNA substrate containing a 23-nt poly(A) tract linked to a 30-nt poly(T). Upon addition of dATP, we observed an upward shift in product mobility (Fig. 4h), indicating extension of the poly(A) using the poly(T) segment as the template. Together, these results reveal that the RT-generated poly(T) functions not only as a primer but also as a template for PP-mediated poly(A) extension.

### Iterative handoff between RT and PP generates palindromic poly(A)/poly(T)-rich duplex-like DNA

Sequencing revealed multiple alternating poly(A) and poly(T) tracts within DRT7 reaction products (Fig. 2e), indicating that DNA synthesis proceeds through iterative cycles rather than a single extension event. Notably, addition of dATP alone produced only a modest increase in product length, whereas inclusion of both dATP and dTTP yielded substantially longer DNA products (Fig. 4h), suggesting that dTTP incorporation can occur following poly(A) synthesis. To directly test whether the RT domain can extend poly(A) with dTTP, we supplied an exogenous 5′ FAM-labeled 120-nt poly(A) oligonucleotide and assayed dTTP incorporation. DRT7 efficiently extended poly(A) with dTTP, and this activity was abolished by RT mutation (Fig. 4i), demonstrating that the RT domain catalyzes dTTP addition onto poly(A) substrates. These results indicate that RT can regenerate poly(T) tracts following PP-mediated poly(A) synthesis.

We next asked how the final palindromic poly(A)/poly(T)-rich duplex-like ssDNA is generated. Our results support a model in which the newly synthesized poly(T)–poly(A)–poly(T) intermediate is returned to the PP domain, where the poly(T) segment again serves as both primer and template for further extension. Iterative cycles of RT-mediated poly(T) synthesis and PP-mediated DNA extension produce molecules containing multiple alternating poly(A) and poly(T) tracts (Fig. 2e). Continued RT–PP cycling progressively elongates the DNA substrate, with the final PP-mediated step reusing previously synthesized segments as templates to generate long, protein-primed palindromic poly(A)/poly(T)-rich ssDNA, consistent with SMRT sequencing data showing complementary poly(A)/poly(T) sequences arranged in a palindromic architecture (Fig. 2e). Together, these findings support an iterative handoff mechanism in which RT and PP cooperate through iterative handoff, with RT serving as both primer and template to repeatedly regenerate poly(T) tracts, and PP repeatedly extending the DNA substrate over successive cycles, ultimately producing long, protein-primed, protein-templated, palindromic poly(A)/poly(T)-rich duplex-like ssDNA.

## Discussion

In this study, we demonstrate that DRT7 confers broad-spectrum resistance through an abortive infection strategy. DRT7 encodes a multifunctional enzyme containing both a RT and a PP domain, and limits phage propagation by synthesizing long, protein-primed, protein-templated, sequence-specific, palindromic poly(A)/poly(T)-rich duplex-like ssDNA through iterative handoff between the RT and PP domains, revealing a distinct strategy of RT-based immunity and an unexpected pathway for nucleic acid synthesis.

In the absence of phage infection, DRT7 predominantly exists as a closed dimer, producing short protein-primed, protein-templated, sequence-specific ∼10-nt poly(dT) intermediates. Phage infection, likely mediated by transcriptional regulators, triggers a transition to an open, active dimer through extension of the RT-synthesized poly(T) strand. Protomer 1 acts as a structural scaffold, whereas protomer 2 undergoes large conformational rearrangements that enable coordinated RT and primase–polymerase (PP) activities. The RT domain initiates protein-primed, protein-templated, thymidine-specific poly(T) synthesis via an arginine-rich recognition pocket. The elongated poly(T) strand is transferred to the PP domain, where it serves as both primer and template for poly(A) synthesis. Then the 3′ end of poly(T)-poly(A) intermediate is returned to the RT domain for renewed poly(T) addition. Iterative RT–PP cycling culminates in a final PP-mediated extension that reuses previously synthesized poly(T)/poly(A) intermediates to produce long, protein-primed, protein-templated palindromic poly(A)/poly(T)-rich duplex-like ssDNA (Fig. 4j). This mechanism shows that DRT7 functions as a protein self-templating DNA polymerization system, which reveals a new pathway for sequence-specific nucleic acid synthesis.

Although the precise biological role of these DNA products remains to be established, their conserved palindromic architecture and strong A/T bias point to DNA structure—rather than primary sequence—underline DRT7 activity (Fig. 2e). The self-complementary structure form stable secondary structures resembling double-stranded DNA yet lack free ends, potentially interfering with phage DNA replication or transcription. Another remaining question is how phage infection activates DRT7. We identified a putative transcriptional regulator that activates DRT7 and triggers cell death *in vivo* (Fig. 1h); however, this factor alone does not stimulate DNA synthesis *in vitro*, suggesting that additional components or cellular contexts may be required for full activation. Finally, although the structural basis for DNA transfer between RT and PP active sites is not yet resolved, coordinated catalytic activities, together with recent preprint evidence of PP-mediated error-prone synthesis coupled to RT proofreading, suggest frequent substrate exchange between the two domains^33^.

In summary, we uncover a distinct RT-based anti-phage defense in which coordinated RT and primase–polymerase activities generate protein-primed, protein-templated, palindromic, self-complementary poly(A)/poly(T)-rich duplex-like ssDNA through iterative synthesis, representing a protein self-templating DNA polymerization system capable of de novo generating palindromic duplex-like DNA structures. These findings expand the functional scope of nucleic acid synthesis pathways and establish a foundation for harnessing DRT-like enzymes in programmable DNA synthesis.

## Supporting information

Cryo-EM table

## Methods

### Plasmid and *E. coli* strain construction

*Escherichia coli* DH5α, BL21 Star (DE3) and DH10B cells were used for plasmid construction, protein expression and phage related experiments, respectively. Phages T4, T5 and T7 were gifts from Xiao Yi’s lab at Shenzhen Institutes of Advanced Technology, Chinese Academy of Sciences. Phages T1(11303-B1), T2 (11303-B2) and T6 (11303-B6) were obtained from the American Type Culture Collection (ATCC). Phage lambda (DSM 4499) and isolates from the·BASEL collection were acquired from the Leibniz Institute DSMZ-German Collection of Microorganisms and Cell Cultures.

### Plasmid cloning, protein expression and purification

The *Escherichia coli* SC366 *drt7* gene, fused to a C-terminal Strep-tag and including a 310-bp upstream region was synthesized by Sangon Biotech and cloned into the pRSFDuet-1 vector. The resulting construct (pRSFDuet-DRT7-Strep) was introduced into *E. coli* BL21 Star (DE3). Protein expression was induced by adding 0.5 mM isopropyl-β-D-thiogalactoside (IPTG, Sangon) when the OD_600_ reached ∼0.8. Following induction, cultures were incubated at 18□ for 20 hours. Cells were then collected by centrifugation and resuspended in lysis buffer consisting of 20 mM Tris-HCl (pH 8.0), 300 mM NaCl, and 2 mM DTT.

For DRT7 purification, cell pellets were disrupted and lysed by AH-1500 High Pressure Homogeniser (ATS, inc.) at 800 bar for 10 min and centrifuged for 30 min at 32,914 × g. The supernatant was loaded onto a Strep-tag II column (Smart Lifesciences). The column was washed five times with lysis buffer. Proteins were eluted with buffer A (20 mM Tris-HCl pH 8.0, 100 mM NaCl, 2 mM DTT) supplemented with 20 mM D-biotin and further purified on a Superdex 200 Increase 10/300 GL column (Cytiva) in a buffer containing 20 mM Tris-HCl pH 8.0, 300 mM NaCl and 2 mM DTT. All mutants were generated by site-directed mutagenesis and purified following the same procedure.

### Phage plaque assays

The *Escherichia coli* SC366 DRT7, along with its upstream 310 bp sequence was cloned into the pACYC184 plasmid. *E. coli* DH10B was transformed with pACYC184 -DRT7 plasmid or an empty pACYC184 vector and cultured on LB agar plates at 37□ in the presence of 10 μg/mL Tetracycline. Single colonies were inoculated in liquid LB media containing the appropriate antibiotic and grown overnight at 37□ with shaking at 200 rpm. On the following day, 500 μL of the overnight culture was mixed with 25 mL of molten MMB top agar (LB supplemented with 0.1 mM MnCl□, 5 mM MgCl□, 5 mM CaCl□, 0.5% agar) and poured onto square plates (Biosharp). The plates were allowed to dry for 1 hour at room temperature. Tenfold serial dilutions of each phage were prepared in phosphate-buffered saline (PBS), and 10 μL of each dilution was spotted onto the bacterial layer. Plates were incubated at 37□ overnight. Plaque-forming units (PFUs) were quantified by counting plaques after incubation. Phage defense activity was evaluated by calculating the fold reduction in efficiency of plating (EOP), defined as the ratio of the plaque-forming units per mL (PFU mL^−1^) obtained on a lawn of empty vector (EV) control cells to the PFU mL^−1^ obtained on a lawn of defense system-expressing cells. All mutants were generated by site-directed mutagenesis and analyzed using the same procedure.

### Phage-infection dynamics in liquid medium

Plasmid pACYC184-DRT7 or a negative control (pACYC184 empty vector) were transformed into *E. coli* DH10B competent cells, respectively. Single colonies were selected and grown overnight in LB medium supplemented with 10 μg/mL tetracycline at 37□ with shaking at 200 rpm. Overnight cultures were diluted 1:100 into fresh LB medium containing 10 μg/mL tetracycline and grown at 37□ until the OD_600_ reached 0.2. 180□µL of the culture were transferred into wells in a 96-well plate containing 20 µL of phage lysate for a final multiplicity of infection (MOI) of 5, 0.5, 0.05 as an uninfected control (MOI=0), 20 μL of PBS was added instead of phage lysate. Optical density measurements at a wavelength of 600 nm were taken every 5 min using Synergy H1 microplate reader (Agilent BioTek) at 37□. Data analysis was performed using GraphPad Prism 9.

### Isolation of escape phage mutants

To isolate phage mutants capable of escaping DRT7-mediated defense, tenfold serial dilutions of phages were plated on *E. coli* expressing DRT7. Plates infected with Bas63 phages were incubated overnight at 37 □, and individual plaques were picked into 3 mL cultures of DRT7-expressing bacteria. Cultures were incubated overnight at 37 □ with shaking at 200 r.p.m., followed by centrifugation at 16,000 g for 5 min to collect phage-containing supernatants. To test the phages for the ability to escape from DRT7 defense, a small drop plaque assay was used as described above. Phages able to escape DRT7 defense were subject to two consecutive rounds of single plaque picking and purification to confirming their escape ability.

### Extraction and sequencing of phage genome

Phage genomic DNA was isolated from 1 mL amplified lysates of mutant and wild-type (WT) phages using a phage DNA isolation kit (Norgen Biotek). Host nucleic acids were degraded by treatment with DNase I and RNase A (1 μg/mL each; TIANGEN) at 37 □ for 30 min, followed by DNase I inactivation at 75 □ for 5 min. Protein digestion was performed by adding Proteinase K (5 μL, 20 mg/mL) and incubated at 55 □ for 30 min.

Subsequently, lysis buffer B (1 mL; provided in the kit) was added, and samples were vortexed and incubated at 65 □ for 30 min. Isopropanol (320 μL) was then added, and the mixture was briefly vortexed and applied to the purification column. After centrifugation at 6,000 g for 1 min, the column was washed three times with 400 μL wash buffer, and the phage genome DNA was eluted in 30 μL double distilled water.

Genomic re-sequencing and mutation analysis were conducted by Iotabiome Biotechnology Co,. Ltd. (Suzhou, China) on BGI DNBSEQ-T7 sequencer.

### Cryo-EM sample preparation and data acquisition

Aliquots (3.5 μL) of purified DRT7 samples (∼15 mg/mL) were incubated with 1mM dATP & dTTP on ice for 5 min before sample preparation and were applied onto the glow-discharged grids (UltrAuFoil 300 mesh R1.2/1.3, Quantifoil). The grids were then blotted for 2.5 s and plunge-frozen in liquid ethane vitrified by liquid nitrogen using Vitrobot Mark IV (FEI Company) at 8□ and 100% humidity. For data acquisition, the grids were transferred to FEI Titan Krios electron microscope operating at 300 kV, and movies (32 frames, total accumulated dose 50 e^−^/Å^2^) were collected using a direct electron detector Gatan K3 in the super-resolution mode with a defocus range from −1.5 to −2.5 μm. Automated single-particle data acquisition was performed with the EPU (Thermo Fisher Scientific) program at a nominal magnification of 105,000, yielding a final pixel size of 0.827 Å.

### Cryo-EM data processing

Micrographs processing was performed by RELION 3.1^34^ and cryoSPARC v4.7.1^35^. Images were first imported into RELION 3.1 and motion correction was performed with MotionCor2^36^. Contrast transfer function (CTF) parameters were then estimated by CTFFind4^37^. Particles were auto picked using the Laplacian-of-Gaussian method, and these extracted particles were subject to several rounds of 2D classification using cryoSPARC v4.7.1^35^. A total of 77,565 particles were used for Ab-Initio reconstruction for DRT7^dATP & dTTP^ complex. After several iterations of heterogeneous refinement, particles corresponding to the DRT7^dATP & dTTP^ complex underwent CTF refinement followed by another round of NU-Refine and subjected to DeepEMhancer^38^ or EMReady2^39^ to reduce noise levels and obtain more detailed experimental maps. In order to improve the density of the PP domain and nascent poly(T)/poly(A) ssDNA, a custom mask was created around the area of interest followed by re-local the map, then a local refinement job was performed. All cryo-EM reconstructions were validated with the gold standard Fourier shell correlation using the 0.143 threshold^35^. Local resolution estimates were calculated from two half data maps in cryoSPARC v4.7.1^35^. Details related to data processing were shown in table S1.

### Atomic model building and refinement

With the assistance of the structural models predicted by AF2^40^, we manually built the atomic models of DRT7^dATP & dTTP^ complex interactively in COOT^41^. Real-space refinement in PHENIX^42^ was used to refine all models against the cryo-EM maps by applying geometric and secondary structure restraints. All structure figures were prepared in PyMol (http://www.pymol.org) and ChimeraX^43^.

### Mass photometry

Mass photometry measurements were conducted at room temperature using a Refeyn OneMP instrument. Microscope coverslips (24 × 50 mm) were cleaned by sonication for 20 min in isopropanol, followed by 20 min in ultrapure water, and dried under a stream of nitrogen gas. A silicone CultureWell™ gasket was placed onto each coverslip to define sample chambers. After applying a drop of immersion oil to the objective, 10 µL of DRT7 sample (∼1 mg/mL) was loaded into a chamber. Data acquisition and initial analysis were performed using the manufacturer’s software with default settings. Mass distributions were processed and visualized in DiscoverMP (Refeyn Ltd.) and plotted as molar mass histograms.

### *In vitro* DNA synthesis assays

Reactions were performed by adding 400 nM DRT7 or its mutants, and the indicated concentration of deoxynucleotides, and either the random primer or no primer, into reaction buffer (20 mM Tris-HCl pH 8.0, 100 mM NaCl, 5 mM MgCl_2_) to a final volume of 100 μL at 37□ for 120 min except for specific designation. Reactions were terminated by the addition of 4 mg/mL Proteinase K (TIANGEN) and 5 mM EDTA, followed by incubation at 65□ for 30 min.

To extract the DNA, the protease-treated solution was heated to 95□ for 10 min to inactivate protease K, cDNA was precipitated by mixing with equal volume of pre-cooled isopropanol and incubated for 1 hour at −80□ or overnight at 4□, followed by centrifugation at 21,130 × g for 15 min. The supernatant was discarded, and the precipitate was washed twice with 75% ethanol solution and left to dry. Finally, the samples were dissolved in 30 μL ddH_2_O with 20 μg/mL RNase A (TIANGEN). cDNA products were run on 1% agarose gel in 1 × tris-acetate-EDTA buffer then stained with GelRed (Tsingke Biotechnology Co., Ltd.) and SYBR Gold (Thermo Fisher Scientific), and visualized with a molecular imager ChempGel 7000 (SageCreation Beijing China).

### Sequencing of cDNA products

For *in vitro* cDNA production, reactions were performed by incubating 400nM of DRT7 WT or DRT7 YVAA mutant with 800 µM dNTP mixtures and 5 mM MgCl_2_ at 37□ for 120 min. cDNA was extracted as previously described.

For *in vivo* cDNA production, overnight cultures of *E. coli* DH10B strains containing pACYC-DRT7 with Flag-tag at C-terminal (DRT7-Flag) or pACYC-DRT7-YVAA mutant (YVAA-Flag) were back-diluted in 200□mL LB and grown at 37□□ to an OD_600_ of 0.8. The cultures were infected with T4 phage at an MOI of 10 and incubated for 30 minutes at 37□ with shaking at 200 rpm. 200 mL cultures were pelleted by centrifugation at 7,500 g for 5□min and washed twice by PBS buffer. The pellet was resuspended in 5 mL of binding buffer (0.1% SDS, 1% Triton X-100, 2 mM EDTA, 20 mM Tris-HCl pH 8.1, 150 mM NaCl). The cells were crosslinked by a UV light crosslinker at the energy of 120 mJ/cm^2^ for 5 min. Cells were lysed by sonication (3s on, 3 s off, 20 min) using an Avestin EmulsiFlex-C3 homogenizer. The lysate was centrifuged at 20,000 × g for 10 min at 4□. The supernatant was incubated with pre-washed Anti-DYKDDDDK (FLAG) Affinity Beads and incubated overnight. Beads were collected by centrifugation at 5,000 × g for 1 min, and the supernatant was discarded. Beads were washed with 0.5 mL wash buffer by gentle resuspension, followed by centrifugation at 5,000 × g for 1 min. The supernatant was removed, and the washing step was repeated three times to eliminate non-specific binding. Bound proteins and nucleic acids were eluted by adding 100 μL acidic elution buffer (0.1M glycine HCl, pH3.0) and incubating at room temperature for 20 min. Beads were pelleted by centrifugation at 5,000 × g for 1 min, and the supernatant was carefully collected without disturbing the beads. Eluted fractions were immediately neutralized by the addition of 20 μL 1M Tris 8.0. DNA was extracted as previously described.

All purified products were sequenced using the Illumina NovaSeq6000 platform at GENEWIZ, Inc. (Suzhou, China). For Long-read sequencing of *in vitro* cDNA products, the purified products were sequenced on the PacBio Sequel platform at GENEWIZ, Inc. (Suzhou, China).Reads were quality-trimmed, and sequences were mapped to the *E. coli* genome, T4 phage genome (for *in vivo* samples), expression plasmid, and cDNA references using BWA.

### Southern blot analysis

DNA samples collected for gel analysis (as described previously) were also used for Southern blotting. Equal volumes (10 μL) of cDNA were loaded onto a 1% agarose gel and run at 120 V. After electrophoresis, the agarose gel was treated with denature buffer (0.5 M NaOH and 1.5 M NaCl) for 20 min. DNA was transferred to a positively charged nylon membrane for 60 min at 260 mA in 0.5 × TBE transfer buffer. The DNA was crosslinked to the membrane using a UV light crosslinker. The membrane was then pre-hybridized in BeyoHybrid™ Hybridization Buffer (Beyotime) for 2 hours at 42□. A biotinylated oligonucleotide probe specific to the target sequence was added to the hybridization buffer at a final concentration of 10 pmol/mL, and hybridization was carried out overnight at 42□.

Following hybridization, membranes were washed twice in low-stringency wash solution (Beyotime) for 5 min each at room temperature, followed by a high-stringency wash in high-stringency wash solution (Beyotime) for 5 min at 42□. Membranes were subsequently washed once with 1× washing buffer (Beyotime) and blocked in blocking buffer (Beyotime) for 15 min at room temperature. Blocking buffer supplemented with streptavidin–HRP conjugate (1:2000 dilution) was then applied, and membranes were incubated at room temperature for 15 min. After incubation, the membrane was washed quickly in 1 × Washing Buffer (Beyotime) for 1 min, followed by four additional washes with 1 × Washing Buffer (Beyotime) for 5 min each. Membranes were transferred to equilibration buffer (Beyotime) and incubated at room temperature for 5 min. The membrane was developed using BeyoECL Moon Working Solution (Beyotime) and imaged using the ChempGel 7000 Molecular Imager (SageCreation, Beijing, China).

### ssDNA extension assay

Reactions were performed by incubating 400 nM DRT7 or its mutants with 800 μM appropriate dNTP mixtures and 100 nM 5′-FAM-labeled ssDNA in reaction buffer containing 20 mM Tris-HCl (pH 8.0), 100 mM NaCl, and 5 mM MgCl□, with the total reaction volume adjusted to 50 μL. For reactions using 5′-6-FAM–labeled 30T20A ssDNA, the substrate was pre-denatured at 95□ for 15 min and cooled to room temperature prior to use. Samples were incubated at 37□ for 60 min in the dark. Reactions were terminated by treatment with Proteinase K for 20 min, followed by heating at 95□ for 5 min. Reaction products were resolved on 10% denaturing urea-polyacrylamide gels in 0.5 × TBE buffer and visualized using a ChemiGel 7000 imaging system (SageCreation, China). The first lane contained FAM-labeled ssDNA incubated with EDTA as a negative control, while the remaining lanes corresponded to protein-incubated reaction products. All assays were repeated at least three times, and representative results are shown.

### Reporting summary

Further information on research design is available in the Nature Portfolio Reporting Summary linked to this article

## Data availability

The cryo-EM density maps have been deposited in the Electron Microscopy Data Bank (EMDB) under accession number EMD-69078 (DRT7 in a closed dimeric state) and EMD-69079 (DRT7 in an open dimeric state). The atomic coordinates have been deposited in the Protein Data Bank (PDB) with accession number 23LM (DRT7 in a closed dimeric state) and 23LN (DRT7 in an open dimeric state).

## Acknowledgments

N. J. is an investigator of SUSTech Institute for Biological Electron Microscopy. We thank the staff at Southern University of Science and Technology (SUSTech) Cryo-EM Center for assistance in data collection on the SUSTech Titan KRIOS cryo-electron microscope. N.J was supported by Guangdong Major Project of Basic Research (grant no. 2025B0303000005); National Natural Science Foundation of China (Grant No. 32570044); Guangdong Natural Science Foundation (Grant No. 2024A1515010541). X.Y.S was supported by National Natural Science Foundation of China (Grant No.325B2001)

## Author contributions

Conceptualization: NJ; *In vivo* and *in vitro* biochemical experiments: XYS, YX, HQ, LL; Cryo-EM sample preparation; data collection, processing and analysis; structure refinement: JTZ, XYW; Writing – original draft: NJ, JTZ; Writing – review & editing: NJ, JTZ, XYS, YX, XYW, HQ

## Competing interests

The authors declare no competing interests.

---

**Extended Data Fig. 1.**
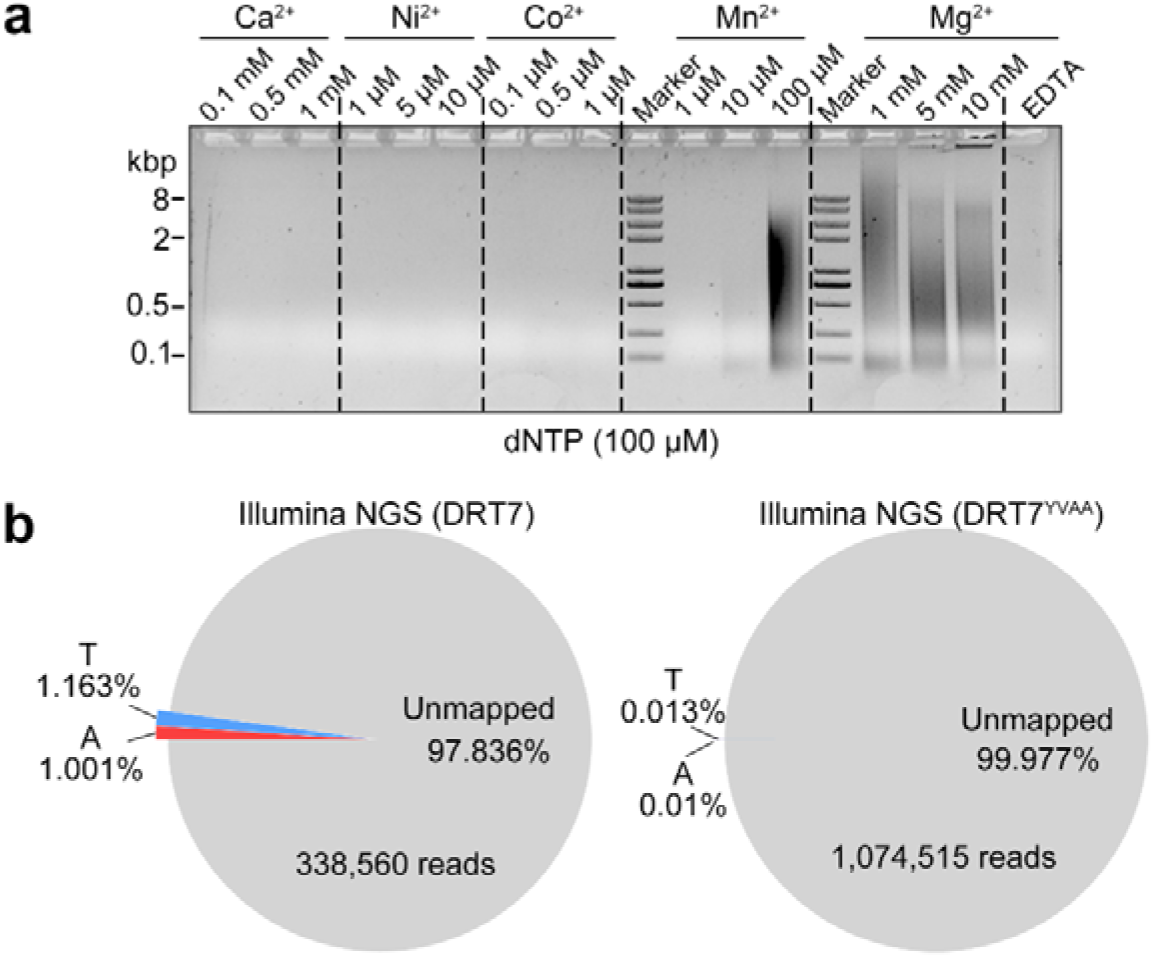
DRT7 produces A/T-rich DNA *in vitro* and *in vivo*. **a**, Divalent cation requirement for DRT7-mediated DNA synthesis. Reactions contained 400 nM DRT7 and 100 μM dNTPs. **b**, Proportion of next-generation sequencing (NGS) reads containing ≥15 consecutive adenine or thymine bases and >60% total A/T content from DNA associated with wild-type DRT7 or a catalytically inactive mutant (D626A/D627A; YVDD→YVAA) in T4-infected *E. coli* DH10B cells.

**Extended Data Fig. 2.**
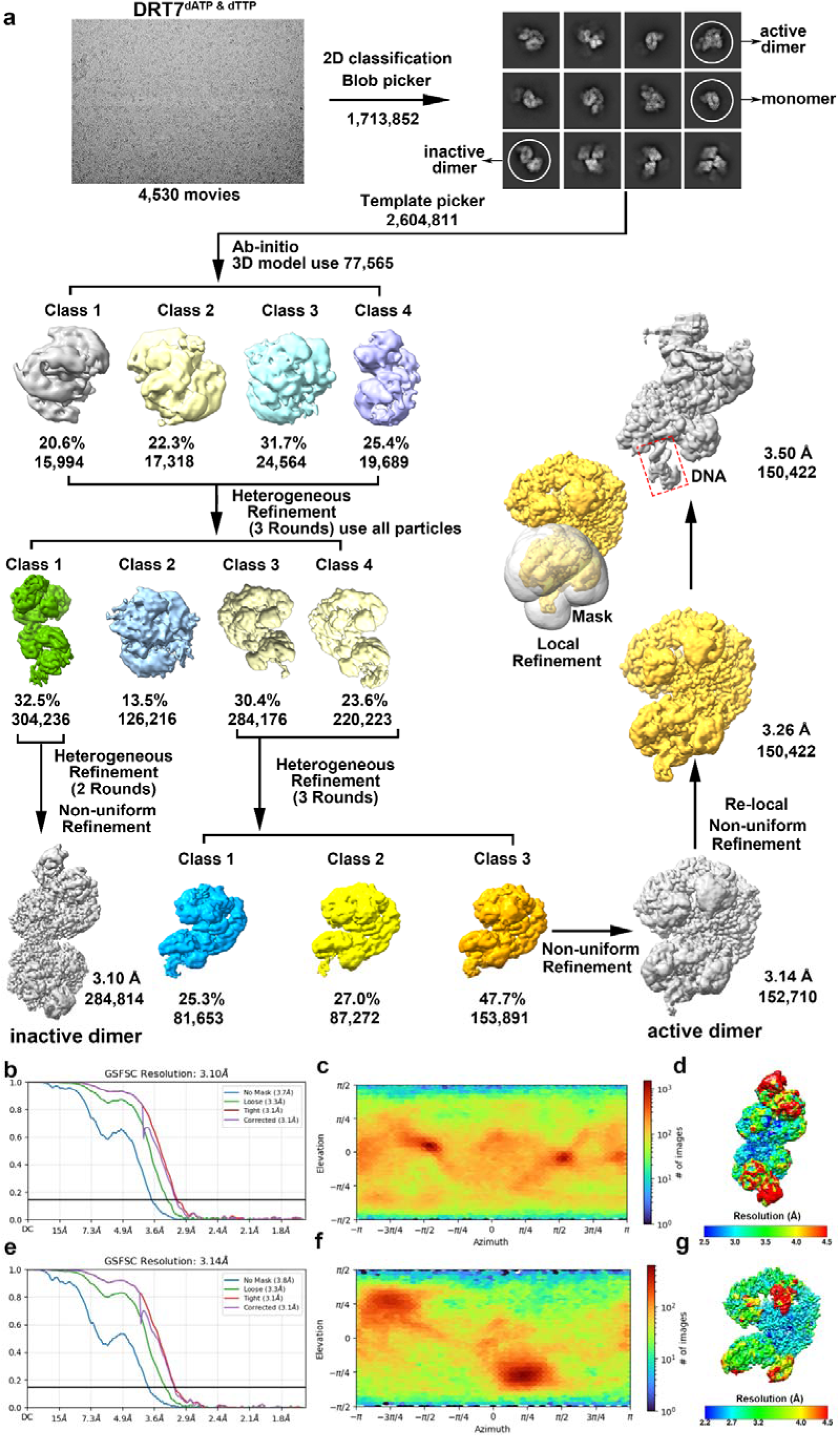
Cryo–electron microscopy reconstruction of DRT7 in the presence of dATP and dTTP. **a**, Image-processing workflow for cryo-EM analysis of DRT7 in the presence of dATP and dTTP. **b**, **e**, Fourier shell correlation (FSC) curves for the inactive (**b**) and active (**e**) states. **c**, **f**, Particle orientation distributions for the inactive (**c**) and active (**f**) states. **d**, **g**, Final 3D reconstructions of the inactive (**d**) and active (**g**) states, colored by local resolution.

**Extended Data Fig. 3.**
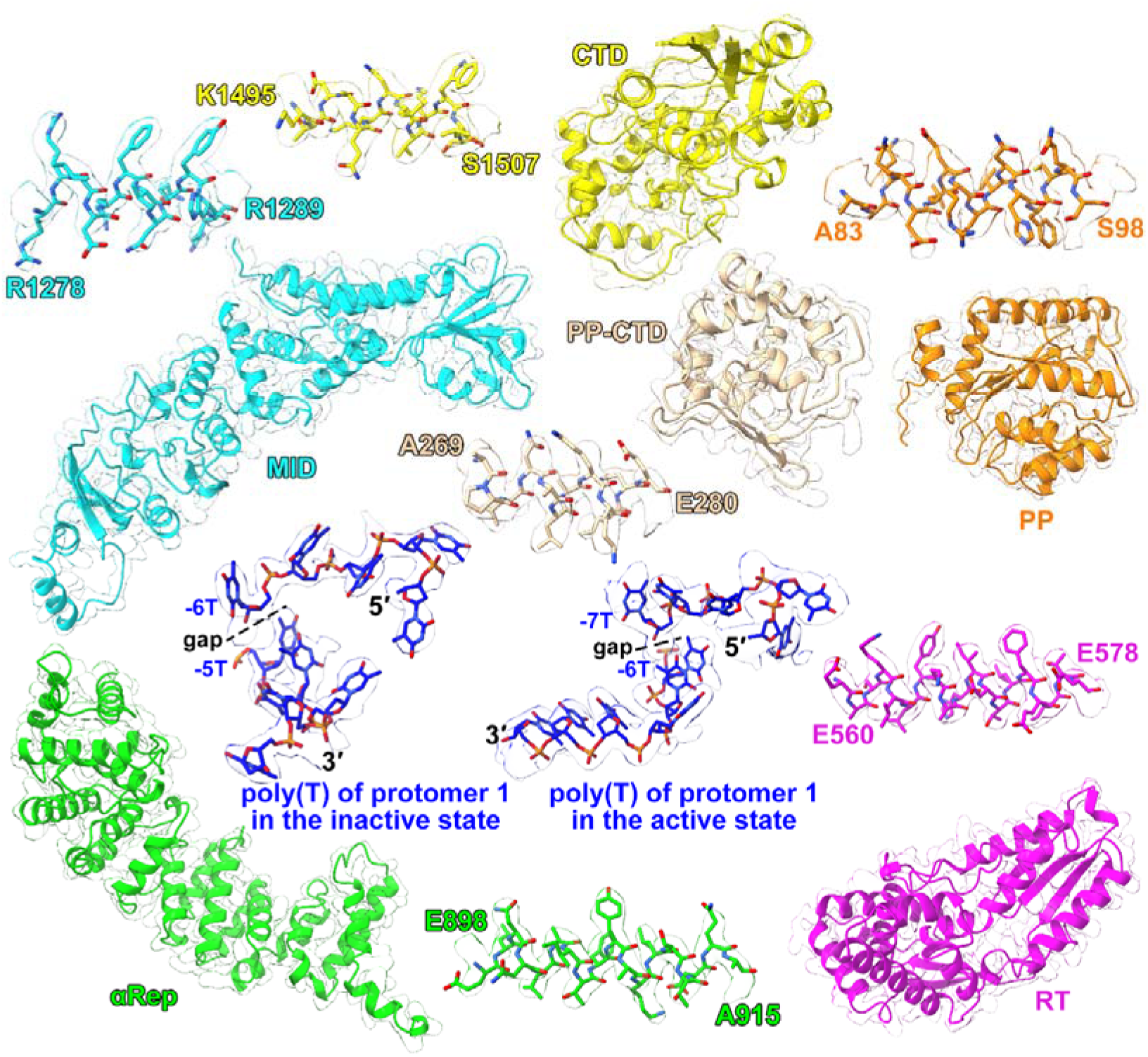
Cryo–EM density map of DRT7 in the presence of dATP and dTTP. Density maps for selected regions are shown in the context of the corresponding atomic models.

**Extended Data Fig. 4.**
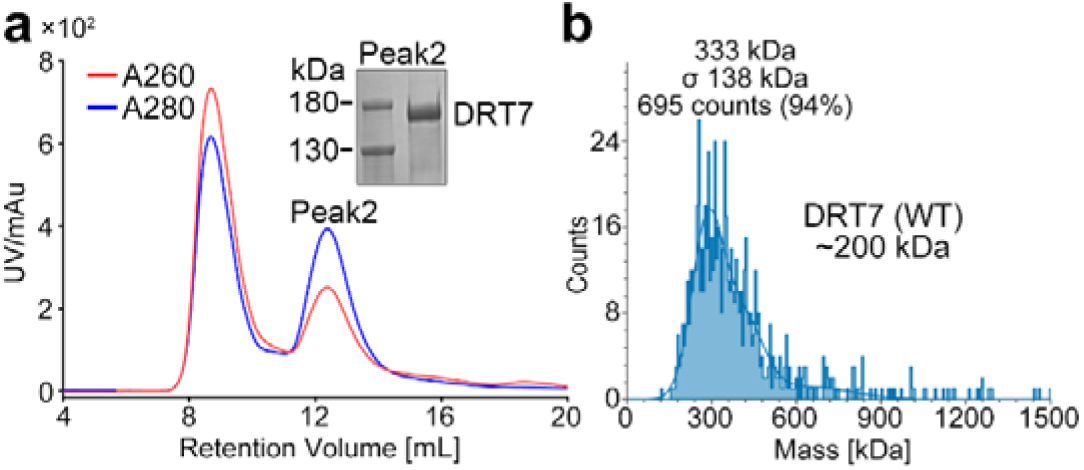
Biochemical characterization of the DRT7 oligomeric state. **a**, Size-exclusion chromatography profile of DRT7. Inset shows SDS–PAGE analysis of fractions corresponding to peak 2. **b**, Mass photometry analysis of DRT7 showing measured molecular masses and relative particle abundances.

**Extended Data Fig. 5.**
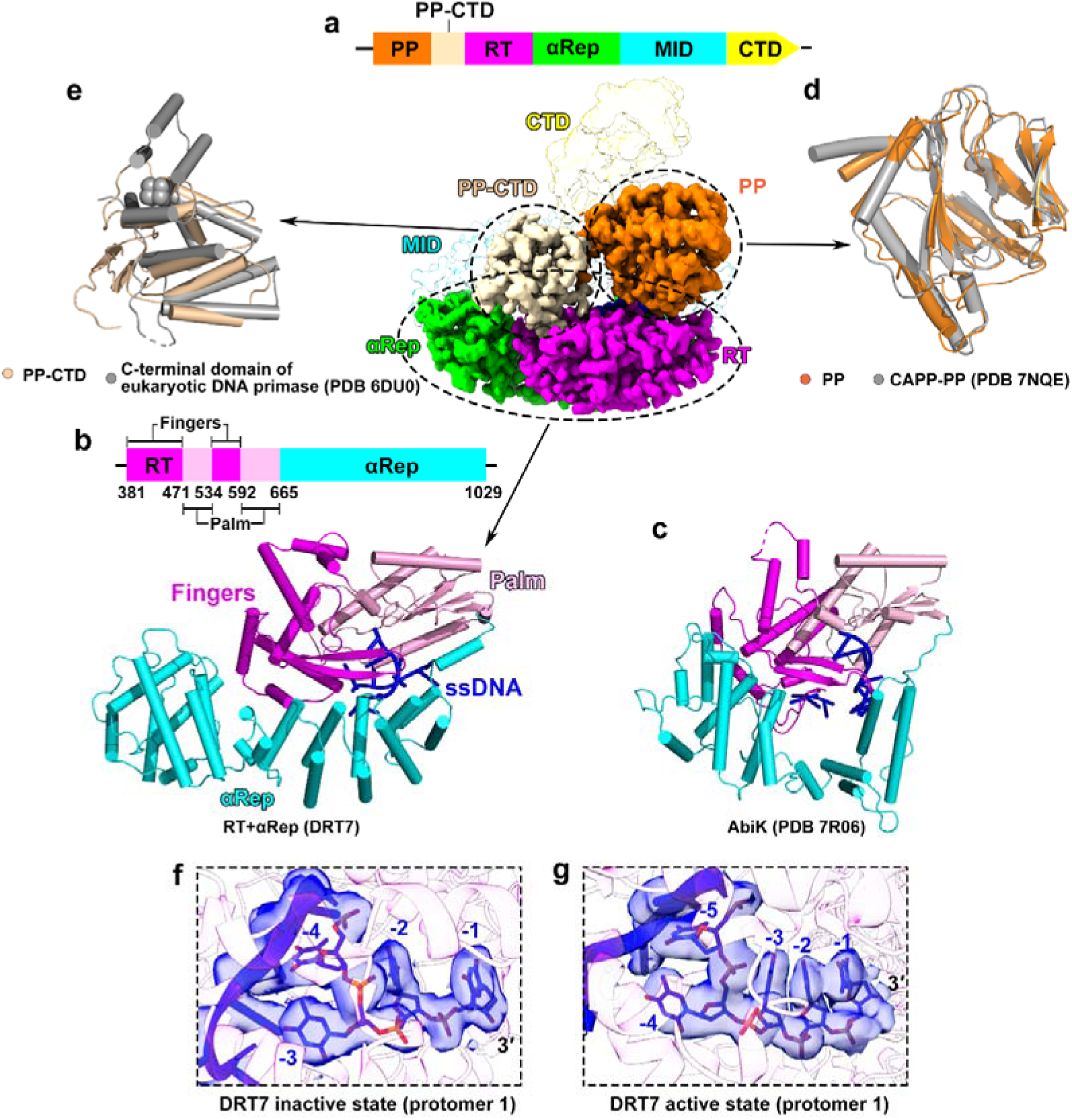
Structural comparisons of DRT7 with related RT and primase–polymerase. **a**, Domain organization of DRT7. **b, c,** Structural analysis of the RT+αRep part of DRT7 (**a**) and AbiK (PDB 7R06) (**c**). **d**, Structural alignment of the primase–polymerase (PP) domain of DRT7 with the PP domain of CRISPR-associated Prim–Pol (CAPP; PDB 7NQE). **e**, Structural alignment of the C-terminal domain (CTD) of the DRT7 PP domain with the CTD of eukaryotic DNA primase (PDB 6DU0). **f**, **g**, Close-up views of the 3′ terminus of ssDNA bound in the RT active site of protomer 1 in the inactive (**f**) and active (**g**) states.

**Extended Data Fig. 6.**
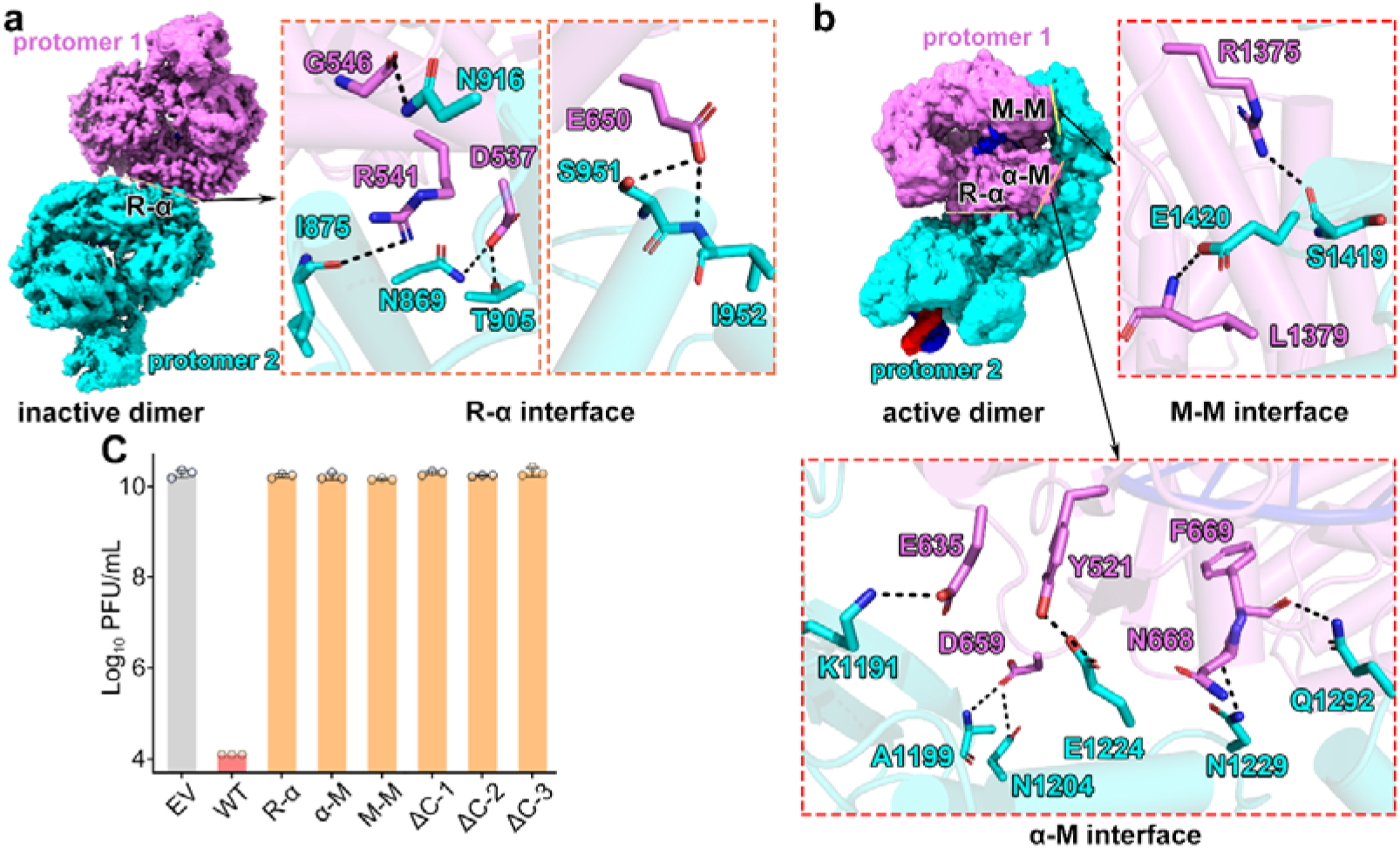
Three dimer interfaces are essential for DRT7 function. **a**, The conserved RT–αRep (R–α) interface is present in both inactive and active states. Insets show detailed interactions at the interface. **b**, The αRep–MID (α–M) and MID–MID (M–M) interfaces are formed specifically in the active state. Insets highlight key intermolecular contacts. **c**, Anti-phage activity of wild-type DRT7 and interface mutants. *E. coli* DH10B cells expressing wild-type DRT7; R–α interface mutants (D537A/K540A/R541A/K563A/K567A/D571A); α–M mutants (E1224A/N1229A/R1298A/Q1292A); M–M mutants (Y1373A/R1375A/N1386A); C-terminal truncations (Δ1038–1776, Δ1122–1776, Δ1458–1776); or empty vector (EV) were challenged with T4 phage. Data represent mean ± SD from three biological replicates with individual data points shown.

**Extended Data Fig. 7.**
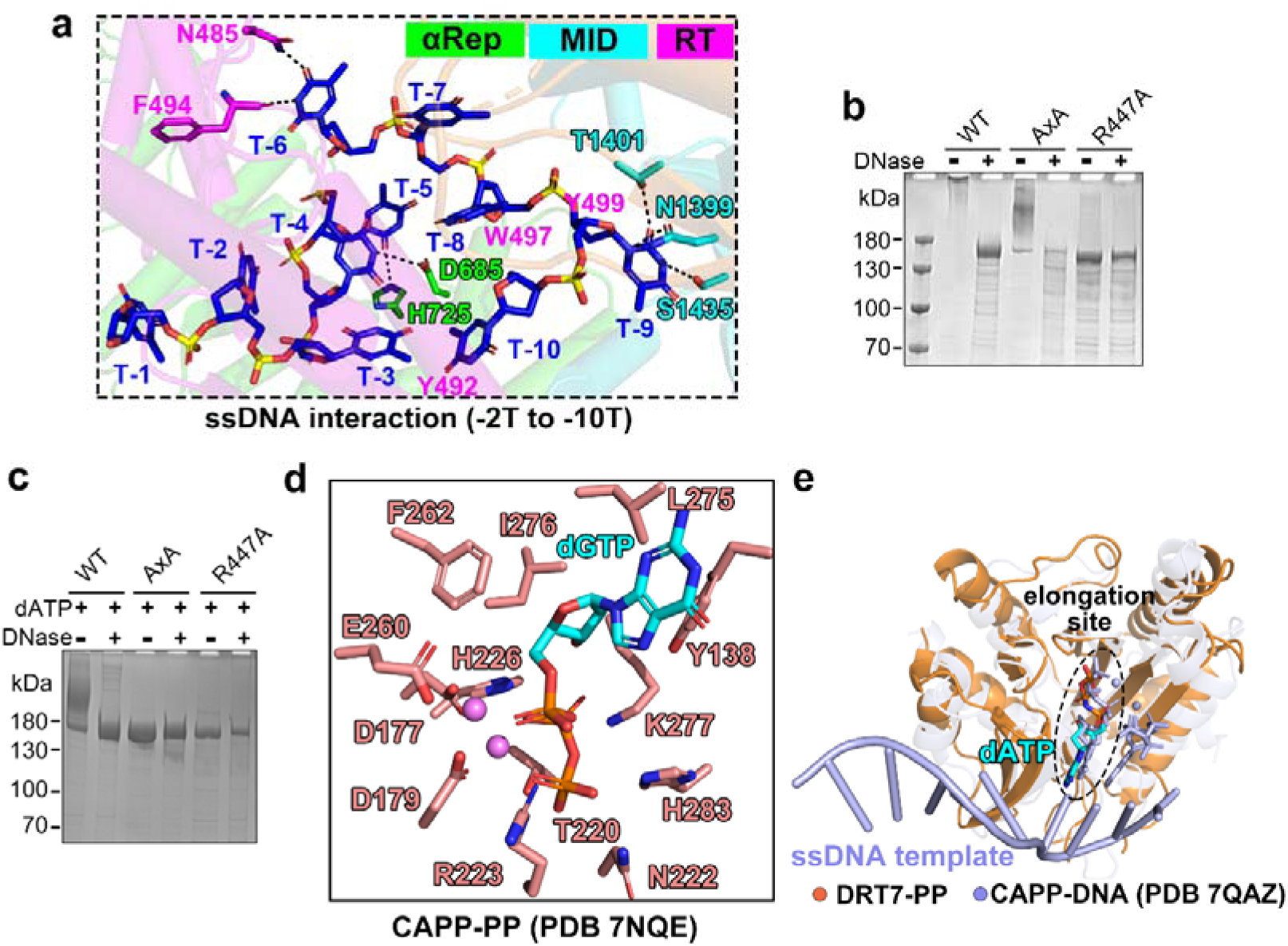
Structural features underlying poly(T) and poly(A) synthesis by the RT and PP domains. **a**, Detailed interactions between poly(T) ssDNA and DRT7, with residues colored by domain. **b**, SDS–PAGE analysis of wild-type DRT7, a PP-deficient mutant (D626A/D627A), and an RT-deficient mutant (R447A) in the presence of 800 μM dNTP. The upward mobility shift caused by covalent DNA attachment is reversed by DNase treatment. **c**, SDS–PAGE analysis of wild-type DRT7, a PP-deficient mutant (D626A/D627A), and an RT-deficient mutant (R447A) in the presence of 800 μM dATP. The upward mobility shift caused by covalent DNA attachment is reversed by DNase treatment. **d**, Close-up view of the active site of the CAPP primase–polymerase domain bound to dGTP (PDB 7NQE). **e**, Structural superposition of the ATP-bound DRT7 PP domain with the CAPP–DNA complex (PDB 7QAZ). ATP from DRT7 is shown in cyan, whereas the ssDNA and paired dNTP from the CAPP complex are shown in light blue.

## Notes

### Competing Interest Statement

The authors have declared no competing interest.

