## Supplementary material for "Antiviral reverse transcriptase–primase synthesizes protein-templated DNA": Cryo-EM table

**Extended Data Table 1| Cryo-EM data collection, refinement and validation statistics**

|  | DRT7 in a closed dimeric state | DRT7 in an open dimeric state |
| --- | --- | --- |
| **Data collection and processing** |  |  |
| Magnification | 105,000 | 105,000 |
| Voltage (keV) | 300 | 300 |
| Electron exposure (e^–^/Å^2^) | 50 | 50 |
| Defocus range (μm) | 1.5 to 2.5 | 1.5 to 2.5 |
| Pixel size (Å) | 0.827 | 0.827 |
| Symmetry imposed | C1 | C1 |
| Initial particle images (no.) | 2,604,811 | 2,604,811 |
| Final particle images (no.) | 150,422 | 284,814 |
| Map resolution (Å) | 3.14 | 3.10 |
| FSC threshold | 0.143 | 0.143 |
| Map resolution range (Å) | 1.6-999 | 2.2-999 |
| **Refinement** |  |  |
| Initial model used (PDB code) | Alaphfold | Alaphfold |
| Model resolution (Å) | 2.10 | 1.99 |
| FSC threshold | 0.5 | 0.5 |
| Map sharpening *B* factor (Å^2^) | -84.1 | -99.7 |
| Map Correlation Coefficient | 0.81 | 0.85 |
| **Model composition** |  |  |
| Non-hydrogen atoms | 21,593 | 28,718 |
| Protein residues | 2,613 | 3524 |
| Nucleotides | 24 | 20 |
| ***B* factor (Å^2^)** |  |  |
| Protein | 88.64 | 86.30 |
| Nucleotides | 32.72 | 47.97 |
| **R.m.s. deviations** |  |  |
| Bond lengths (Å) | 0.015 | 0.014 |
| Bond angles (°) | 1.414 | 1.164 |
| **Validation** |  |  |
| MolProbity score | 2.38 | 1.94 |
| Clash score | 17.25 | 11.22 |
| Poor rotamers (%) | 2.79 | 1.93 |
| **Ramachandran plot** |  |  |
| Favored (%) | 95.61 | 97.14 |
| Allowed (%) | 4.39 | 2.86 |
| Disallowed (%) | 0.00 | 0.00 |
| **EMDB** | EMD-69078 | EMD-69079 |
| **PDB** | 23LM | 23LN |
